# Increased oscillatory power in a computational model of the olfactory bulb due to synaptic degeneration

**DOI:** 10.1101/2020.08.06.239293

**Authors:** J. Kendall Berry, Daniel Cox

## Abstract

Several neurodegenerative diseases impact the olfactory system, and in particular the olfactory bulb, early in disease progression. One mechanism by which damage occurs is via synaptic dysfunction. Here, we implement a computational model of the olfactory bulb and investigate the effect of weakened connection weights on network oscillatory behavior. Olfactory bulb network activity can be modeled by a system of equations that describes a set of coupled nonlinear oscillators. In this modeling framework, we propagate damage to synaptic weights using several strategies, varying from localized to global. Damage propagated in a dispersed or spreading manner leads to greater oscillatory power at moderate levels of damage. This increase arises from a higher average level of mitral cell activity due to a shift in the balance between excitation and inhibition. That this shift leads to greater oscillations critically depends on the nonlinearity of the activation function. Linearized analysis of the network dynamics predicts when this shift leads to loss of oscillatory activity. We thus demonstrate one potential mechanism involved in the increased gamma oscillations seen in some animal models of Alzheimer’s disease and highlight the potential that pathological olfactory bulb behavior presents as an early biomarker of disease.

## I. INTRODUCTION

The olfactory system, and in particular the olfactory bulb (OB), is implicated in early stages of a number of neurodegenerative diseases, including two of the most prevalent, Alzheimer’s disease (AD) and Parkinson’s disease (PD) [1–3]. In both AD and PD, olfactory deficits occur years before diagnosis and often before other symptoms [2, 4–9]. Furthermore, the OB is a site of early pathology in both diseases [7, 10–15], with resulting aberrant neural activity [16–21]. We hope that computationally modeling olfactory bulb activity in disease-like conditions can help to further shed light on mechanisms of dysfunction, identify markers of disease, and bring attention to the opportunity the OB presents for earlier diagnosis of neurodegenerative illnesses.

The OB is the first processing area for incoming odor information [22], but exactly how it is represented is an ongoing question for which there are various theories, mainly revolving around combinatorics of principal neuron activity [23–26]. Oscillations in neural activity in the bulb may also play a part in encoding odor identity, and are likely important to odor recognition or information transfer, or both [27]. Neural networks can exhibit a variety of dynamical behaviors [28]; the push-pull nature of the excitatory-inhibitory interactions of the OB make it an interesting example of a nonlinear oscillator [29]. In addition, studying the robustness of the system’s oscillations has relevance to the neurodegenerative diseases mentioned above.

The oscillatory behavior of the OB in the gamma range (40-80 Hz) is driven by reciprocal synaptic interactions between the dendrites of excitatory mitral cells and inhibitory granule cells [27, 30] (also called dendrodendritic synapses [31]). That is, mitral cells excite granule cells which in turn inhibit the mitral cells, leading to gamma band oscillatory activity. Other frequencies of oscillations are present in the bulb as well, namely theta (2-12 Hz) and beta (15-30 Hz). The precise manner in which PD or AD impacts these oscillations and other OB functions is still a matter of investigation [1, 2], although studies have found perturbations in this oscillatory activity in animal OBs in the presence of both PD-like pathology [16, 32] and AD-like pathology [18–21, 33].

More generally, the effects of PD and AD pathology on neurons is a very active area of study, with various alterations of neuron function resulting from overexpression or injection of pathological protein (see [34, 35] for reviews of PD pathology, see [36–38] for reviews of AD pathology). Synaptic dysfunction is one effect for which there is evidence in both AD [39, 40] and PD [41–44], with studies finding (for example) decreased spine density [17, 19, 45], decreased synaptic proteins [19, 20], reduced vesicle release [46], increased synaptic junction distance [20], and decreased synaptic transmission [47].

Computational studies of the effects of AD on neural networks have focused on largely on the hippocampus and cortical areas, especially effects on memory formation and storage (see [48] for review). In PD, most computational models simulate various effects of dopamine loss in the basal ganglia network, exploring changes in network output and oscillatory activity, as well as effects of deep brain stimulation therapy (see [49, 50] for reviews). Importantly, we are not aware of any works that examine the impact of neurodegenerative damage on a computational model of the olfactory bulb.

Many excellent and insightful computational models of the OB exist [51], focusing on various aspects of the olfactory system, such as generating oscillatory behavior [52–54], glomerular layer computations [55], and odor computations and representation [25, 56].

In the present study, we implemented the Li-Hopfield model [52], a rate based model with units representing small populations of neurons. The model replicates gamma band oscillatory activity found in the OB (see [52] compared to electroencephalogram (EEG) studies [57] and micro-electrode recordings of extra-cellular potential, also called local field potentials [58]). The oscillations are produced solely via interaction between model mitral cell (MC) units and granule cell (GC) units, as supported by findings that dendrodendritic interactions between MCs and GCs alone were sufficient for gamma band oscillatory activity [58, 59]. While the model excludes certain aspects of the OB network (such as beta oscillations [60]), it captures some key behaviors, and its simplicity enables semi-analytical and numerical treatments that are not accessible in more complex models.

The original Li-Hopfield work only implemented a 1D connection architecture; here, we expand the model to various sizes and 2D connection structures. We found that the 1D and 2D networks operate similarly, but in different regimes. Importantly, on the scale of the original Li-Hopfield model, we found that for some types of damage, 2D networks show a significant enhancement of gamma oscillatory power at moderate levels of damage to the connections between MCs and GCs. Analysis of the model network’s behavior shows that this results from an increased excitability of the MC population due to a reduction of inhibition, with the nonlinearity of the activation function being essential. The balance of excitation and inhibition is important for robustness of oscillatory activity. As a result, we would expect to see an increase in oscillatory power at moderate levels of damage to the reciprocal connections between excitatory and inhibitory cells in these types of oscillatory networks.

For the remainder of this paper, we first detail the governing equations of the Li-Hopfield model and describe how the model is modified for larger sizes and 2D connection structure. We then lay out the method of delivering damage to the network and explain how oscillatory activity in the network is characterized. The resulting increase in oscillatory activity and underlying mechanisms are explored, with observations about the differences between 1D and 2D networks. Finally, we discuss the relevance to experimental studies, the limitations, and the future directions of the work presented here.

## II. METHODS

### Li-Hopfield Model

The Li-Hopfield model describes the internal state (representative of membrane potential) and output state (or cell activity, representative of firing rate) of mitral and granule cells over an inhale-exhale cycle. Each MC and GC model unit represents a subset or small population of MCs or GCs, with the weight matrices *H*_0_ and *W*_0_ representing the dendro-dendritic synaptic connections between MCs and GCs. For simplicity, MC-MC and GC-GC interactions are not considered here since synaptic interactions in the region giving rise to gamma oscillations are predominantly reciprocal MC-GC synapses [30, 58, 59].

The model is governed by the following set of equations,

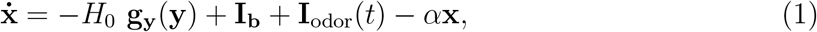

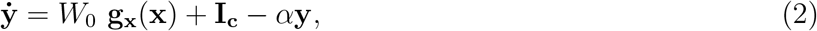

where **x** and **y** are vectors containing the internal state of each MC unit and each GC unit, respectively. The functions **g_x_** (**x**) and **g_y_** (**y**) are sigmoidal activation functions [52] that translate internal state into output state, **I_b_** is tonic uniform background excitatory input to the mitral cells, **I_c_** is tonic uniform excitatory centrifugal input to the granule cells, and *α* is the decay constant, which is taken to be the same for mitral and granule cells in this model. Random noise is added to **I_b_** and **I_c_** in the form given in the original Li and Hopfield paper [52]. **I**_odor_ is excitation from the odor input, which rises linearly with inhale and falls exponentially with exhale. In principle, **I**_odor_ could have different levels of input for each mitral cell. In the simulations here, we defined it to be uniform for simplicity and because damage should affect all odors. The exact functions and values for the parameters can be found in Table I, and are as given in [52, 61]. *H*_0_ and *W*_0_ are the matrices that define the connections from granule to mitral cells (inhibitory, as indicated by the negative sign) and from mitral cells to granule cells (excitatory), respectively.

**TABLE I.** The parameters for the model are as given by Li and Hopfield [52, 61]. The model is evaluated at time steps representing 1 ms and runs for 395 ms. Thus all times *t* below are in ms.

| Parameter | Value |
| --- | --- |
| $I_{b,i}$ | 0.243 |
| $I_{c,i}$ | 0.1 |
| $\alpha$ | 0.15 |
| $g_y(y_i)$ | $\begin{cases} 2.86 + 2.86 \tanh(\frac{y_i-1}{2.86}) & \text{if } y_i < 1 \\ 2.86 + .286 \tanh(\frac{y_i-1}{.286}) & \text{if } y_i \geq 1 \end{cases}$ |
| $g_x(x_i)$ | $\begin{cases} 1.43 + 1.43 \tanh(\frac{x_i-1}{1.43}) & \text{if } x_i < 1 \\ 1.43 + .143 \tanh(\frac{x_i-1}{.143}) & \text{if } x_i \geq 1 \end{cases}$ |
| $I_{\text{odor},i}$ | $\begin{cases} 0 & \text{if } t < 25 \\ 0.00429(t - 25) & \text{if } 25 \leq t < 205 \\ 0.00429(t - 25)e^{-0.03(t-205)} & \text{if } t \geq 205 \end{cases}$ |

The weight matrices *H*_0_ and *W*_0_ dictate the connective structure of the network. The original Li-Hopfield model contained 10 MC units and 10 GC units, connected in mitral-granule pairs, with each pair connecting to neighboring pairs on a 1D ring (Fig. 1(a)(b)). For the work here, the network was adapted to include larger numbers of MCs and GCs. This was accomplished by initializing a matrix of the desired size with entries of the same order magnitude as the original connection matrices, and then updating the non-zero entries randomly until the desired behavior was achieved. Because the interface between the dendrites of the MCs and GCs, the external plexiform layer, lies on the surface of an ellipsoid, matrices were constructed with 2D architecture for each size in the same way (Fig. 2). Therefore, the model was implemented in six different architectures: 20 cells (mitral plus granule), 40 cells, and 100 cells in 1D and 2D. It should be noted that *W*_0_ in the original Li-Hopfield model included extra connections that make the architecture not truly 1D. These connections have been retained for the 1D 20 cell network, but are not present in any of the other 1D network structures. The specific matrices for each size can be found on the Github repository (see Code Accessibility).

**FIG. 1.**
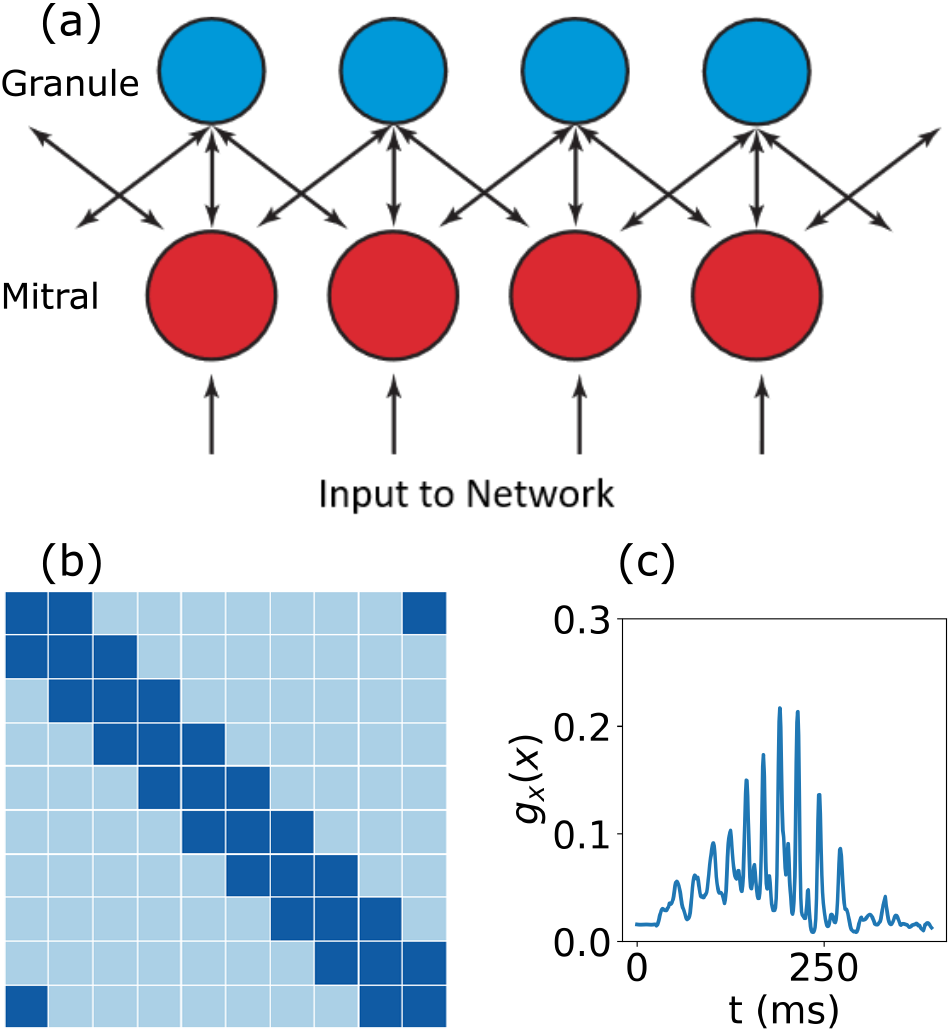
Li-Hopfield Model. (a) Mitral cell units receive odor input and excite the granule cell units, which in turn inhibit the mitral cell layer. The connections between mitral and granule cell units define a 1D ring structure. The networks implemented here have 20 units (mitral plus granule), 40 units, and 100 units. (b) Example weight matrix defining a 1D periodic network. Light blue entries are zero (no connection), dark blue signifies positive non-zero entries (established connection). (c) Example of mitral cell output (*g_x_*(*x_i_*)) over the course of a single inhale-exhale cycle. The inhalation peaks at 205 ms, at which point the exhale begins. Parameters for the model are found in Table I. Each mitral or granule cell unit should be considered as representing a particular population of mitral cells.

**FIG. 2.**
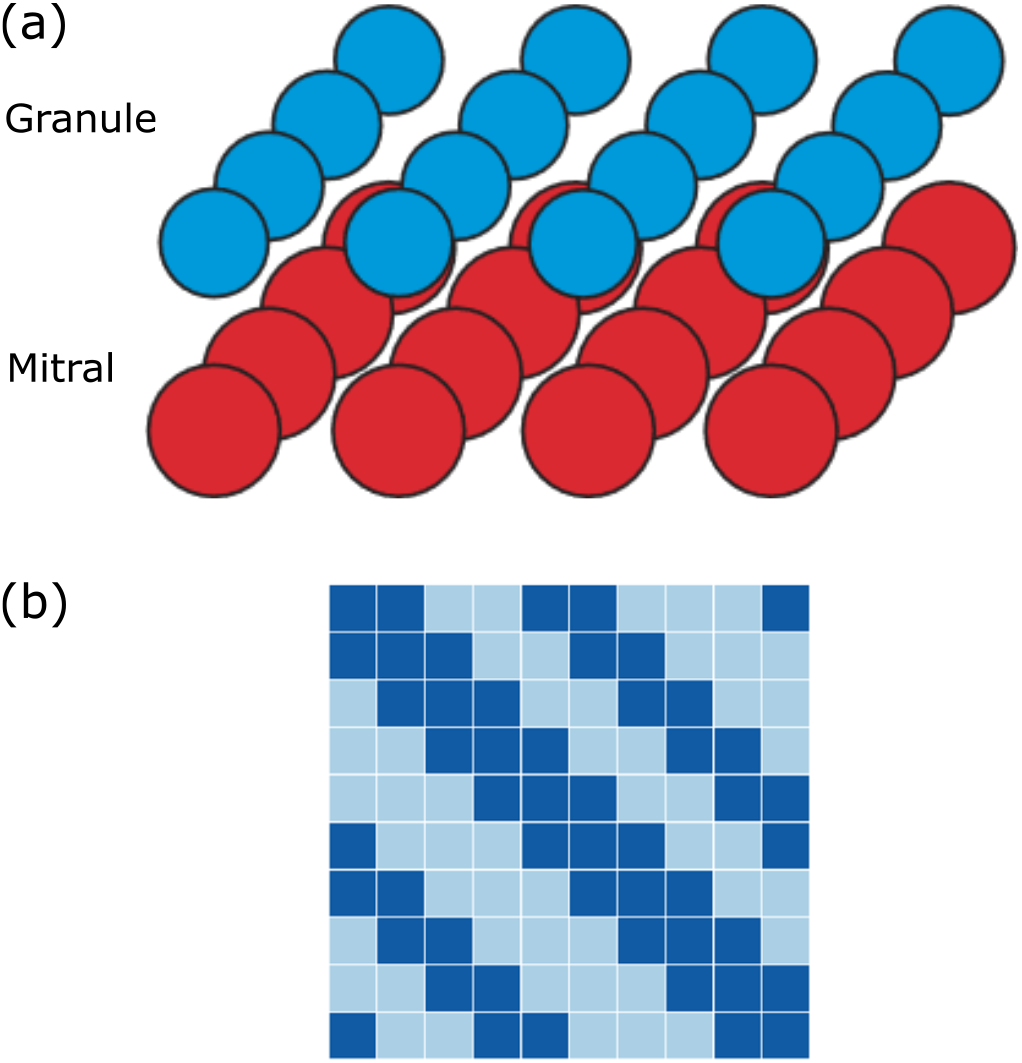
2D Model. (a) The connections between mitral and granule cell units are largely the same as in 1D, but extend in two directions. (b) Example of a weight matrix structure defining a 2D periodic network structure. Light blue entries are zero (no synaptic connection), dark blue signifies positive non-zero entries (established synaptic connection). Each MC unit connects to its GC pair, as well as four other GC units. GCs connect to MC units following the same pattern.

### Damage

In our work here, we focus on damage delivered to the weight matrices *H*_0_ and *W*_0_ (although damaging other network components was also explored, see [62]). This was partly informed by the numerous studies that point to synaptic dysfunction as one salient effect of neurodegenerative pathology [19, 20, 41–44, 46]. Additionally, this aligns with the scope of the Li-Hopfield model, which focuses on the gamma oscillations that arise from the excitatory-inhibitory interactions between MCs and GCs.

We measure damage to the network by *δ*, the fraction of weight removed. In the case of *W*_0_ for example,

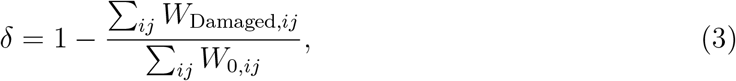

where *W*_0_ is the undamaged matrix and *W*_Damaged_ is the damaged matrix.

In a given trial, damage is delivered to either *H*_0_ (the synaptic connections from granule to mitral cells), or to *W*_0_ (the synaptic connection from mitral to granule cells). The damage to the selected part of the network is increased, the network runs at that damage level, and the activity is recorded. The damage is propagated in one of three ways: Flat Damage (FD), Columnar Damage (CD), or Seeded Damage (SD).

For FD, the damage is delivered to every element of the selected matrix equally. This amounts to simply scaling the chosen quantity uniformly. For example, if FD was applied to *H*_0_, each element of *H*_0_ would be reduced by the same fraction of its original value on each damage step. This continues until the matrix is reduced to zero (see Fig. 3(a)). FD thus represents a global progression of pathology.

**FIG. 3.**
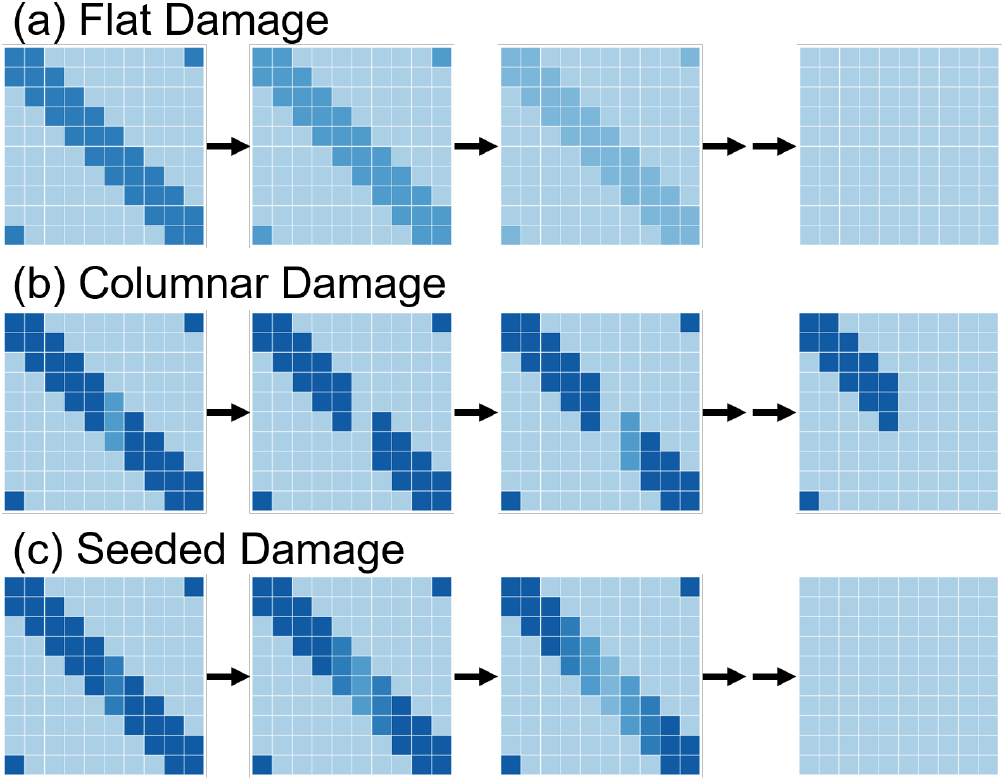
Schematic of Damage Propagation Strategies. (a) Example of FD delivered to *H*_0_ or *W*_0_ in the 1D 20 unit network. (b) Example of CD delivered to *H*_0_ or *W*_0_ in the 1D 20 unit network. Damage begins in a single column (in this case, column 6). For CD, only up to half the matrix weight is removed because in most cases, the network activity was already greatly disrupted by that point. (c) Example of seeded damage delivered to *H*_0_ or *W*_0_ in the 1D 20 unit network. Damage begins in a single column (in this case, column 6).

For CD, the damage is delivered to a specific matrix column (representing synaptic transmission from a particular unit), ramped up until that element is reduced to zero, and then that procedure is repeated successively on adjacent elements until the maximum damage level is reached (see Fig. 3(b)). For example, if CD was delivered to *H*_0_, damage would be delivered incrementally to a single column until it was reduced to zero. The same process would then begin on the column to the right, and this process is continued until half of the columns are removed (Fig. 3 (b)). This is representative of a very local pathological spread.

SD is a hybrid of FD and CD. Damage is first delivered to a single column, and on the next damage step, damage is delivered to that column again, as well as to neighboring elements. For example, if SD was enacted on *H*_0_ in a 1D network, it would begin on one column, say column 6. On the subsequent damage step, the damage would then be delivered to columns 5, 6, and 7. This spreading continues with each damage level until the matrix is reduced to zero (Fig. 3(c)). SD thus represents a pathology that begins locally but becomes more global as damage spreads.

### Characterizing Network Oscillatory Activity

The average oscillatory power, *P*_avg_, is calculated by first high-pass filtering the mitral cell activity above 15 Hz to ignore theta band (2-12 Hz) activity, which was beyond the scope of the present study. Next, the power spectrum (**P** (*f*)) is calculated for each mitral cell from 125 ms to 250 ms using Scipy’s periodogram function [63]. This time window captures the oscillatory behavior during the most active part of the cycle (see Fig. 1(c)) while ignoring the spurious signals that can arise at higher levels of damage that are not actually due to gamma band oscillatory activity (see Fig. S6 [62]). The power spectrum is then integrated over all frequencies, *f*, for each cell (see Fig. S6 through S9 [62] for example power spectra) and averaged over the mitral cell population (*N*) to get *P*_avg_,

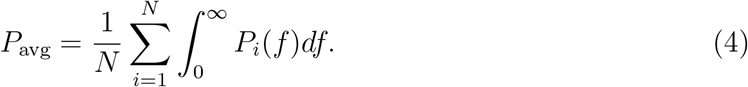

Each quantity is averaged over five trials with differently seeded noise. For CD and SD, the trial is then repeated using each cell as the starting point and the values are averaged again over all starting cells.

### Code Accessibility

The simulations are run in Python using Scipy’s solve ivp function [63] on Ubuntu 18.04, and all code/software described in the paper is freely available online at https://github.com/jkberry07/OB_PD_Model.

## III. RESULTS

### Damage to *W*_0_ and *H*_0_

The effect of damage on average oscillatory power depends on the damage scheme. FD and SD to *H*_0_ or *W*_0_ in 2D networks results in increases in *P*_avg_ at intermediate levels of damage (Fig. 4(a)), but CD rarely shows an increase in *P*_avg_ (Fig. S3 [62]). Thus, an increase of oscillatory power requires global progression of damage to some degree. While this rise in *P*_avg_ is ubiquitous among 2D networks with FD and SD, *P*_avg_ decreases monotonically for most 1D cases (see Fig. 4(b)). For the work shown here, damage was only delivered to either *H*_0_ or *W*_0_, but delivering damage to both has a similar effect of increased oscillations at intermediate levels of damage (see Fig. S5 [62]) and relies on the same principles presented hereafter.

**FIG. 4.**
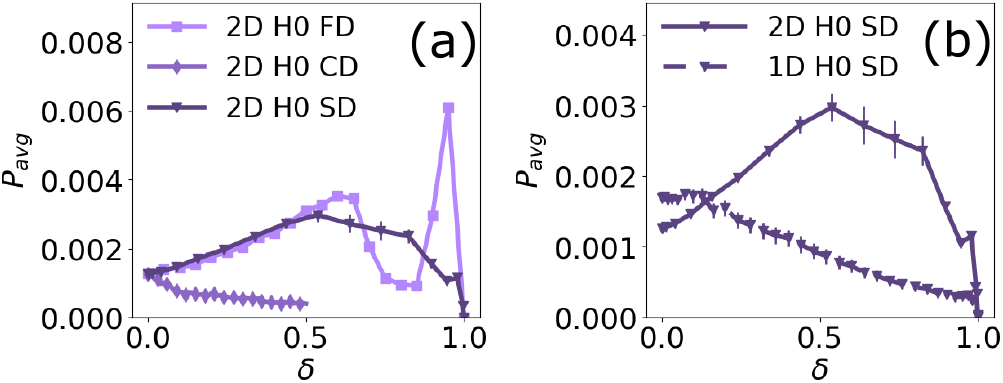
Effect on Oscillatory Power. (a) Average oscillatory power (*P*_avg_) for FD, CD, and SD delivered to *H*_0_ in the 2D 100 cell network plotted against damage as a fraction of total synaptic weight removed (damage level, *δ*). (b) *P*_avg_ for SD delivered to *H*_0_ in the 1D and 2D 100 cell networks. FD and SD to *H*_0_ or *W*_0_ result in a rise in *P*_avg_ for the 2D network, while CD to the 2D network and any kind of damage to the 1D network do not in general (see Figures S2 though S4 [62] for *P*_avg_ for each network and damage type). The sharp peak in *P*_avg_ for FD to *H*_0_ at *δ* = 0.95 is due to a sharp rise and drop in MC unit output states rather than to oscillatory activity, as illustrated by Figure S9 [62]. Example cell activity at various *δ* and associated power spectra can be found in Figures S6 and S7 [62].

The increase in oscillatory power in 2D networks results from larger MC activity amplitude (as opposed to, say, recruitment of previously inactive units). Fig. 5 shows an example cell from the 2D 100 cell network at no damage and at the level of flat damage corresponding to the maximum *P*_avg_. The increase in amplitude shown is seen for all active cells in 2D networks receiving FD. Similar effects are seen for SD, though the following treatment will be done for FD for simplicity.

**FIG. 5.**
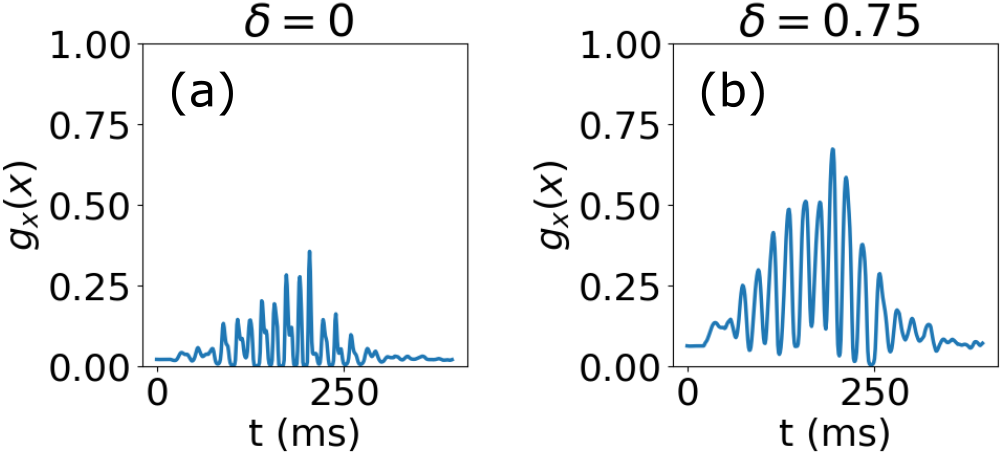
Cell Activity Compared at Different Damage Levels. (a) Output state of mitral cell number 35 in the 2D 100 cell network with no damage. (b) Output state of mitral cell number 35 in the 2D 100 cell network with FD to *W*_0_ at damage level 0.75 (*W*_Damaged_ = 0.75*W*_0_). This was the damage level at which average oscillatory power was maximized.

The question is then why does the output state amplitude increase? To understand this, we look at the internal state of the MC units. The increase in output state amplitude is not due to increased amplitude of internal state oscillations. Rather, it is because of increased average internal state. As a simple measure of internal state amplitude, we inspect the largest peak and the lowest trough between 180 ms and 220 ms (the time window over which when output state oscillations are the greatest) for active units (defined as units that had an individual oscillatory power greater than 0.001). When the null damage case is compared to the damage level at which *P*_avg_ is maximized, the distance from peak to trough is similar (0.7562 *±* 0.0808 for *δ* = 0 and 0.6765 *±* 0.1300 for *δ* at which *P*_avg_ is maximized). The difference is in the increase in the average internal state and how that translates into output state. Though both the minimum trough and the maximum peak increase, only the increase in maximum peak has an effect on the output state because the trough still lies below the threshold of the activation function (see Fig. 6). This illustrates that the nonlinearity is essential for the increase in *P*_avg_ to manifest.

**FIG. 6.**
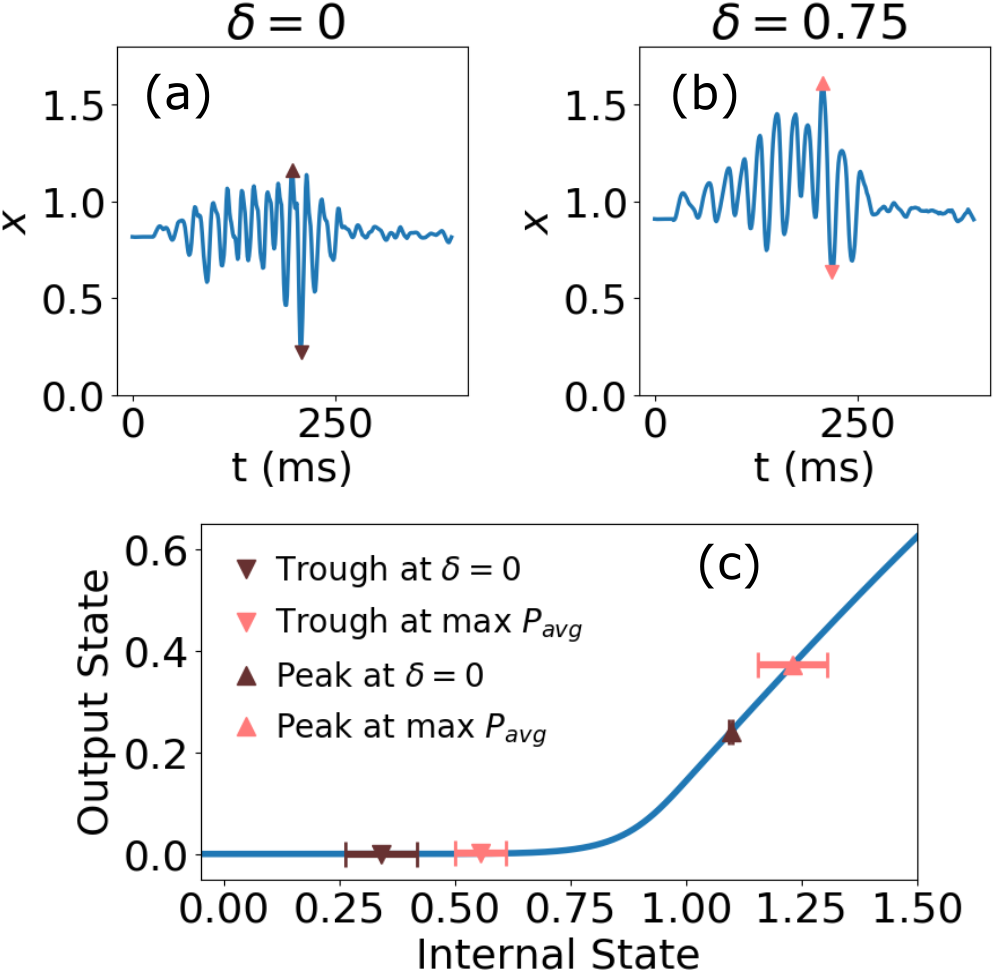
Increase in Average Internal State Results in Increased Amplitude in Output State. (a) Internal state of mitral cell number 35 in the 2D 100 cell network with no damage. (b) Internal state of mitral cell number 35 in the 2D 100 cell network with FD to *W*_0_ at damage level 0.75 (*W*_Damaged_ = 0.75*W*_0_). This was the damage level at which average oscillatory power was maximized for FD to *W*_0_. (c) Activation function *g_x_*(*x*), with dark brown marking the average minimum trough and average maximum peak for 2D networks at *δ* = 0, and light pink marking the same but at the damage level corresponding to the maximum *P*_avg_.

The average internal state increase results from damaging *W*_0_ or *H*_0_, which lowers the overall inhibition to the MC layer, either directly (in the case of *H*_0_) or by reducing excitation to the GC layer (in the case of *W*_0_). The dependence on *H*_0_ and *W*_0_ is explored further in a simplified semi-analytic approach in the supplement [62].

In the Li-Hopfield work, to gain understanding of the full numerical solution, an adiabatic approximation is made in which the oscillations are modeled as variations around a fixed point. This linearized analysis can explain the drop in *P_avg_* that follows the rise. Here, we summarize the analysis done by Li-Hopfield; for a more detailed treatment, see the original work [52]. Keeping the full network, we treat **x** and **y** as deviations from the fixed points, and the governing equations become:

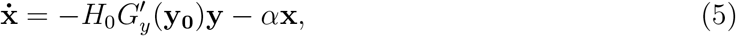

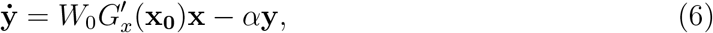

where *G′_y_* (**y_0_**) and *G′_x_*(**x_0_**) are diagonal matrices resulting from the linearized approximation of *g_y_*(*y*) and *g_x_*(*x*). Further manipulation yields:

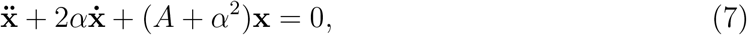

where

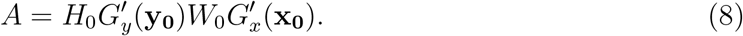

The analytical solution is 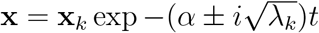, where **x**_*k*_ is the *k^th^* eigenvector of A and *λ_k_* is the *k^th^* eigenvalue. The eigenvalues of A predict the presence of oscillatory behavior.

If

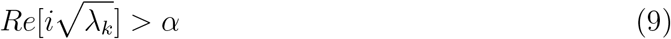

is satisfied, oscillations are present, with the eigenvalue that results in the value highest above *α* dominating [52]. If no eigenvalue satisfies the condition, oscillations die away quickly.

As *W*_0_ or *H*_0_ decrease with damage, the eigenvalues of the matrix A also decrease. Once the dominant eigenvalue falls below the threshold set by the decay rate, the network behaves like a damped oscillator and *P*_avg_ drops dramatically. Though this is derived from an approximation, it faithfully predicts the sudden reduction in *P*_avg_ (see Fig. 7).

**FIG. 7.**
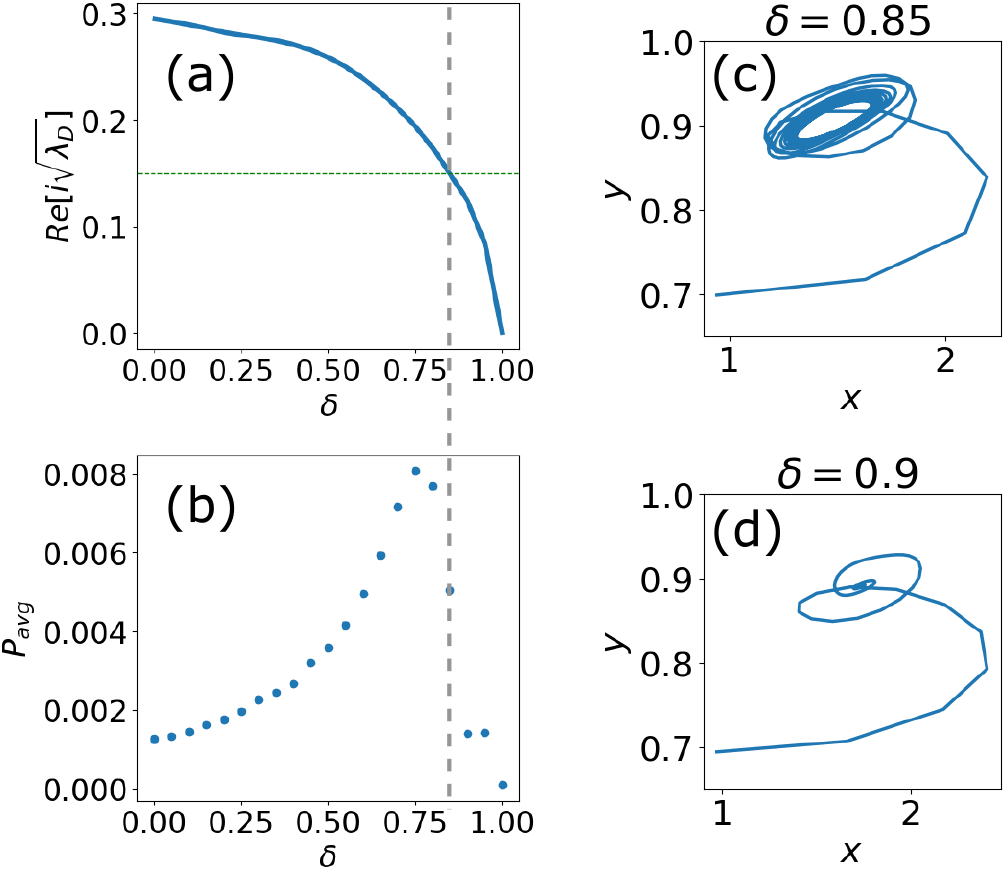
(a) The real part of the square root of the dominant eigenvalue 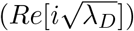 plotted against damage for the 2D 100 cell network with FD to *W*_0_. According to the linearized analysis, when 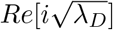 falls below the decay rate, marked as a horizontal dotted green line, oscillations are dampened. (b) Average oscillatory power (*P*_avg_) plotted against damage for the 2D 100 cell network with FD to *W*_0_. The steep drop off in *P*_avg_ corresponds to the damage level at which 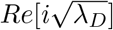 falls below the decay rate, as illustrated by the vertical dashed line. (c) Phase plot, with mitral cell internal state (*x*, mitral cell number 35) on the x-axis, and granule cell internal state (*y*, granule cell number 35) plotted on the y axis, at the damage level immediately before oscillations are quenched. The damage was FD to *W*_0_ in the 2D 100 cell network. The activity settles into a limit cycle, demonstrating oscillatory behavior. (d) Same as (c), but at the following damage level. The activity decays to a fixed point. For the plots in (c) and (d) only, an odor input that is constant in time was used.

This can also be demonstrated with a phase diagram, plotting a MC unit internal state against a GC unit internal state (similar to [29, 64]). To illustrate the network’s behavior in this way, the odor input was modeled as constant in time. Just before the eigenvalue falls below the threshold, the network activity follows a limit cycle (see Fig. 7(c)). However, once it drops below threshold, a bifurcation occurs and network activity approaches a fixed point (see Fig. 7(d)).

### 1D Networks

Network connection matrices of all types were constructed to give roughly equivalent starting oscillatory powers with the same average synaptic weights values. Because the 2D networks were of necessity less sparse than the 1D networks, this meant that the total synaptic weight in the 2D networks was greater than for the 1D networks. Under these conditions, we observe the increase in oscillatory power in FD trials only in 2D networks (for example, Fig. 4). However, if the null-damage matrices in 1D networks instead have a similar total weight rather than the same average weight, the behavior is largely the same, just on a smaller scale.

As an example, we deliver FD to *W*_0_ on a 1D network, but begin with *W*_0_ thrice its typical value. With more weight in the network, the initial average internal state is reduced, and *P*_avg_ begins at a lower value. As damage is applied and the network approaches its null condition (at *δ* = 2/3 because *W*_0_ is tripled), *P*_avg_ increases. This creates a trajectory of behavior with damage that closely resembles that of the 2D network (Fig. 8(a)(b)).

**FIG. 8.**
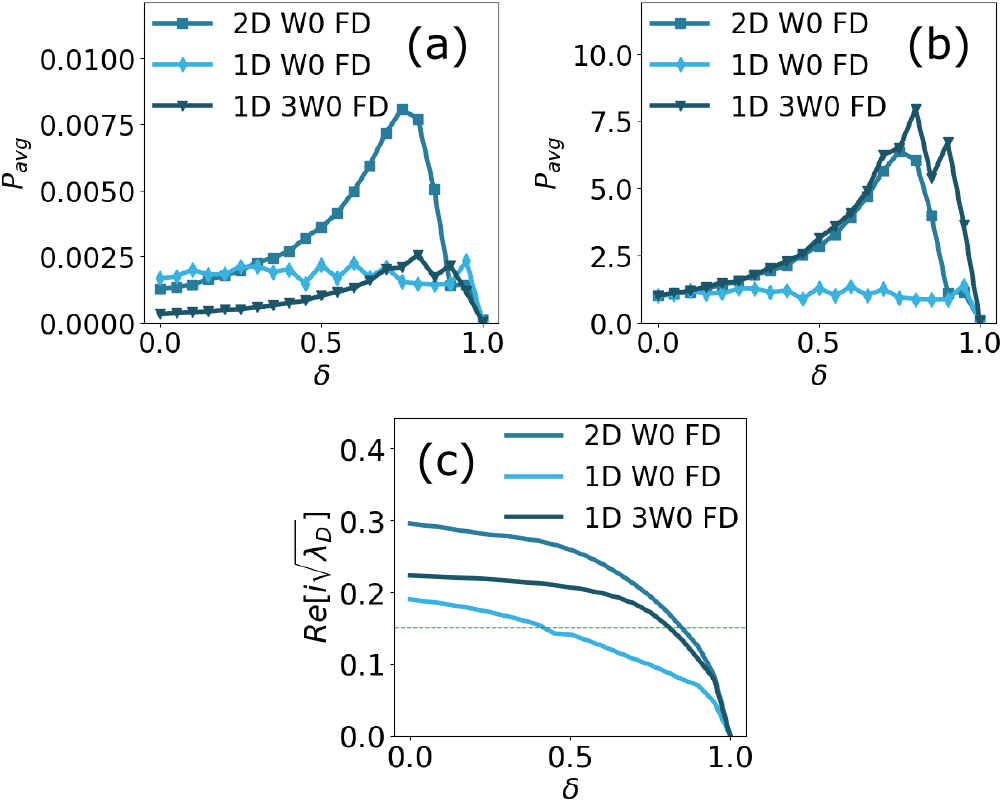
(a) Average oscillatory power (*P*_avg_) for FD delivered to *W*_0_ in the 2D 100 cell network, the 1D 100 cell network, and the modified 1D 100 cell network. The modified network starts the damage trial with *W*_0,modified_ = 3*W*_0,unmodified_. The modified network shows a rise in *P*_avg_, similar in shape to the 2D network. (b) Same as in A, but with *P*_avg_ normalized with respect to its initial value in each case, illustrating more clearly the similarity in behavior between the modified 1D and the 2D networks. (c) The real part of *i* times the square root of the dominant eigenvalue 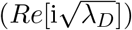 plotted against *δ* for FD delivered to *W*_0_ in the 2D 100 cell network, the 1D 100 cell network, and the modified 1D 100 cell network. The dominant eigenvalue of the modified network still starts below that of the 2D network, but it stays above threshold until a similar level of damage.

This is illustrated also by the behavior of the leading eigenvalue, which indicates the presence of oscillations (as seen in Fig. 7). With a larger weight matrix, the leading eigen-value of the matrix *A* starts at a larger value, and its trajectory is similar to that of the 2D networks. The unmodified 1D network’s leading eigenvalue begins about 2/3 along the trajectory of the modified and so crosses threshold at a lower level of damage (see Fig. 8(c)).

These results demonstrate that the primary difference between the 1D and 2D networks is that they operate in different regimes: the maximum activity state the 2D networks can sustain is greater than that of the 1D networks. Thus, at the initial oscillation size in this model, the 2D networks are below their maximum oscillatory power, while the 1D networks are already at near maximum. The mechanism underlying this difference in maximum activity is beyond the scope of this paper, but it could be due to a greater capacity for cooperative effects resulting from the greater level of connectivity in the 2D networks.

## IV. DISCUSSION

### Seeded Damage

The linearized analysis here is carried out for Flat Damage, but Seeded Damage, which spreads outward from a starting unit, shows a similar increase in oscillatory power at moderate levels of damage and relies on the same mechanisms. That we found similar behavior for SD is important as it better represents a potential method of pathology progression. In PD, for example, misfolded α-synuclein spreads from cell to cell before neuron death [65, 66] in what many believe is a prion-like manner [67–69], although the precise mechanism and the level of damage in the donor cell before transmission is still an active area of investigation [70, 71]. Similarly, evidence suggests prion-like spread of AD pathology as well, of both Aβ [72, 73] and tau [74–76].

### Relation to Aberrant Olfactory Bulb Activity in Animal Models of Disease

Our model’s mechanism for the increase in gamma band oscillatory power is the increase in MC activity. This could be representative of higher firing rates of participating MCs, or of recruitment of less active MCs within the population represented by the active units (as previously inactive units rarely activate in our model, though they theoretically could). The increase in MC activity is due to a reduction of inhibition, either by decreasing GC excitation from the MCs (damage to *W*_0_, which could represent less AMPA receptor activation in GCs) or by decreasing inhibition of MCs by GCs (damage to *H*_0_, which could represent less GABA receptor activation in MCs by GABA receptor loss or reduced GABA transmission from GCs). Either damaging *W*_0_ or *H*_0_ (or both) could be consistent with loss of dendrodendritic synapses. Generally, our model implies that reduced GABA transmission leads to increased gamma oscillations, a result relevant to several experimental studies.

W. Li *et al.* and Chen *et al.* measured OB activity in mice expressing amyloid precursor protein and presenilin 1 (APP/PS1) and found significant increases in gamma band power associated with synaptic deficits [18, 19]. Both groups found that treatment with a GABA agonist decreased the heightened gamma power, suggesting that increases in the gamma band may have been due to a decrease in GABA-induced inhibition to the MCs. W. Li *et al.* also measured increased MC firing rates, although measurements of cell activity were done on OB slices rather than *in vivo* [19]. Note that by including PS1, these two studies may be more specifically relevant to familial AD rather than to sporadic AD [77], and it is not clear that the effects seen were due to Aβ alone. Other studies have also found increased gamma power in OBs of transgenic AD mice models [20, 21], with S. Li *et al.* working with expression of p-tau (P301S mice) rather than APP. While they also found an increase in gamma power and impaired synaptic function, MC firing rates were decreased rather than elevated, suggesting other mechanisms at play. Note also that because AD-like pathology was induced by gene expression in all of these studies [18–21], it is more relevant to the FD modeled here rather than to SD.

The observations of these studies closely relate to work by Lepousez and Lledo [58], who found that GABA antagonist increased oscillations in the gamma band and GABA agonist had the opposite effect. However, they found that MC firing rates were not significantly affected by GABA antagonist (except possibly for increased excitation of MCs that were initially less active), instead showing that the power increase was likely due to increased MC synchronization. Additionally, they found an important reliance on NMDA channels in GCs, which could be relevant to PD since NMDA receptors may be among the targets of pathological α-synuclein [41]. Our model is limited in this regard by a lack of explicit NMDA activity and a clear mechanism for varying synchrony.

While the research mentioned above found increases in gamma oscillations, it should also be noted that some studies have found decreases in OB oscillations in the presence of Aβ. Hernández-Soto *et al.* applied Aβ by injection and measured activity *in vivo* in rat OBs an hour after application [33], and Alvarado-Martínez *et al.* measuring cell activity in mouse and rat OB slices *in vitro* after bath application of Aβ [78]. Both found overall decreases in OB activity.

To our knowledge, few studies have measured olfactory bulb neural activity in the presence of PD-like pathology. Kulkarni *et al.* recorded local field potentials in mice olfactory bulbs after injection of α-synuclein pre-formed fibrils directly into the OB. They measured a significant increase in oscillatory power in the beta band (15-30 Hz) following incubation periods ranging from 1-3 months. Zhang *et al.* modeled PD pathology in mice by reducing the population of dopaminergic neurons in the substantia nigra in mice [32]. They measured an increase in the spontaneous oscillatory power of all bands, theta (2-12 Hz), beta (15-35 Hz), and gamma (36-95 Hz). The full extent of the mechanisms underlying the observations in both of these studies is outside the scope of this model. Dopaminergic input to the OB from the substantia nigra was not modeled here, although it could be possible that the net effect of reducing that input is less inhibition to the MC population. As for the first study, beta oscillations in particular require centrifugal input and rely on channels not modeled here (although the same interactions modeled here are also critically involved, see [60, 79]). The Li-Hopfield model includes centrifugal input to the GCs in only a superficial way, and greater detail is required to reproduce beta oscillations in the OB.

Of note, thus far, experimental studies have shown differences between OB activity in PD-like pathology compared to AD-like pathology. Though modeling these differences lies outside the scope of the present model, if the disease-specific alterations in oscillatory power in the olfactory bulb prove robust, it provides a potential tool for differentiation in early diagnosis of neurodegenerative disease.

## Conclusions and Future Directions

Despite the limitations of the model presented here, we believe it is nevertheless relevant to investigating the effects of neurodegenerative damage on oscillatory activity in the olfactory bulb. The balance between inhibitory and excitatory activity between the MC and GC populations is essential to gamma oscillations and depends on multiple mechanisms and principles [58]. Here, we highlight one principle, which is that as long as inhibition is great enough, marginal decreases in inhibitory action will increase gamma oscillations due to the nonlinear nature of the neural activity [80]. Because it’s unlikely that a biological network would begin close to the regime of maximum oscillatory power (and thus close to the drop off seen in Fig. 7), we would expect to see this effect with moderate damage to the inhibition.

The model here spotlighted one mechanism that may be in play, and serves as a proof of concept that computational modelling can help give insight into OB dysfunction. More detailed models of the OB network and activity are needed to explore other mechanisms and effects of neurodegenerative damage. For example, working with a model based on work by Osinski *et al.* [81] or David *et al.* [82] may offer insight into the modulation of beta oscillations found by Kulkarni et. al [16]. And a model similar to that by Li and Cleland [83] may help investigate effects of cholinergic perturbation in AD, as reviewed by Doty [2].

The prevalence of olfactory dysfunction in neurodegenerative disease presents both an opportunity and a challenge; it is a common early symptom [1–3], yet its applicability in diagnosis is limited by the broadness of its presence. Thus it is important to continue the study of the mechanisms and behavior of OB oscillations in the presence of neurodegenerative damage since, as discussed above, aberrant OB oscillatory activity may show a point of differentiation between PD and AD pathology. Additionally, recent progress in non-invasive measurement of human olfactory bulb activity [84] brings this area of research closer to clinical relevance. For example, Iravani *et al.* used surface electrodes to measure electroencephalogram (EEG) activity originating from the olfactory bulb in human patients, specifically in the gamma range [84]. As these techniques are developed, understanding the aberrant OB activity in neurodegenerative diseases could be a powerful tool for realizing earlier diagnosis of these illnesses.

## ACKNOWLEDGMENTS

We acknowledge useful conversations with M. Goldman, D. Wesson, and A. Kulkarni, and we thank D. Wesson and A. Kulkarni for critical readings of the manuscript. This grew from preliminary work on the problem by Avinash Baidya, and we thank him for useful discussions about the project.

