## Supplemental Material for "Increased oscillatory power in a computational model of the olfactory bulb due to synaptic degeneration"

(Dated: July 2, 2021)

### I. OTHER NETWORK COMPONENTS DAMAGED

Trials were also run with damage delivered to components of the network beside $W_0$ and $H_0$, namely to:

- Granule cell layer
- Mitral cell layer
- $\mathbf{I}_{\text{odor}}$, the input to mitral cells from the glomerular layer

Internal damage to the mitral cell layer (MCL) or to the granule cell layer (GCL) is implemented by multiplying the right-hand-side of the differential equation (except for the leak term) for the given cell by some fraction less than one, $(1 - \delta_i)$. For example,

$$\dot{x}_i = (1 - \delta_i)(-\sum_j H_{0,ij}\ g_{y,j}(y_j) + I_{b,i} + I_{\text{odor},i}(t)) - \alpha\ x_i$$

would be damage delivered to the $i^{th}$ mitral cell unit. In this case, $\delta$ is calculated as

$$\delta = \sum_i \frac{\delta_i}{N},$$

where $N$ is the number of mitral cells.

*

### II. SUPPLEMENTAL FIGURES

**Figure S1: Fixed Point Dependence on $H_0$ and $W_0$**

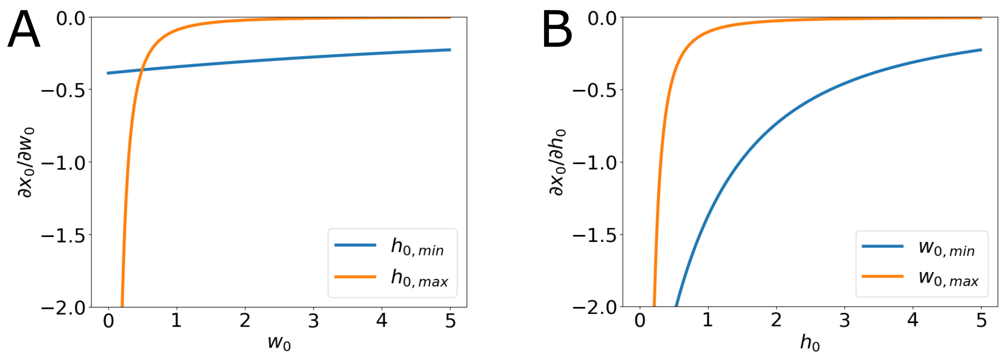


In the Li-Hopfield work, to gain understanding of the full numerical solution, an adiabatic approximation is made in which the oscillations are modeled as variations around a fixed point. Under this simplification, the internal state fixed point is a fair representation of the average internal state. With that assumption, we can explore fixed point dependence on $H_0$ and $W_0$. To make the following analysis tractable, we make two simplifications. First, we reduce the network to a single MC unit and a single GC unit. Second, we approximate the activation functions with a first-order Taylor series expansion around the observed average internal state of active units ($x_{\text{avg,obs}}$ and $y_{\text{avg,obs}}$),

$$\begin{aligned} g_y(y) &\approx g_y(y_{\text{avg,obs}}) + g'_y(y_{\text{avg,obs}})(y_0 - y_{\text{avg,obs}}) \\ &= a_y + b_y y_0, \end{aligned} \tag{1}$$

$$\begin{aligned} g_x(x) &\approx g_x(x_{\text{avg,obs}}) + g'_x(x_{\text{avg,obs}})(x_0 - x_{\text{avg,obs}}) \\ &= a_x + b_x x_0, \end{aligned} \tag{2}$$

where $a_y = -0.6847,\ b_y = 0.9693,\ a_x = -0.2132,\ b_x = 0.8223$ are constants based on the average internal state during trials delivering flat damage to $H_0$ and $W_0$. Thus the fixed point equations become

$$0 = -h_0(a_y + b_y y_0) - \alpha x_0 + I_{\text{odor}} + I_b, \tag{3}$$

$$0 = w_0(a_x + b_x x_0) - \alpha y_0 + I_c. \tag{4}$$

We can now analytically solve for $x_0$ and differentiate with respect to $w_0$ and $h_0$,

$$\frac{\partial x_0}{\partial w_0} = \frac{\alpha h_0 b_x b_y [h_0(I_c b_y/\alpha + a_y) - I_o - \alpha a_x/b_x]}{(h_0 w_0 b_x b_y + \alpha^2)^2}, \tag{5}$$

$$\frac{\partial x_0}{\partial h_0} = \frac{-\alpha^2 b_y [w_0(I_o b_x/\alpha + a_x) + I_c + \alpha a_y/by]}{(h_0 w_0 b_x b_y + \alpha^2)^2}. \tag{6}$$

For simplicity, $I_{\text{odor}}$ is set to a constant value and collected with $I_b$:

$$I_o = I_b + I_{\text{odor}}(t = 175ms) = 0.99375$$

.

When plotted, it becomes apparent that for all values of $h_0$ and $w_0$ within the range used in the work here, the derivatives are negative, meaning that a decrease in $W_0$ or $H_0$ increases $x_0$, as seen in FD to $W_0$ and $H_0$. In A, the partial derivative of the fixed point with respect to $w_0$ is plotted against $w_0$ using the minimum and maximum values of $H_0$ in the 100 cell 2D network ($H_{0,\text{min}} = 0.0017,\ H_{0,\text{max}} = 2.0606$). The plot in B is the partial derivative with respect to $h_0$ plotted against $h_0$ using the minimum and maximum values of $W_0$ in the 100 cell 2D network ($W_{0,\text{min}} = 0.0163,\ W_{0,\text{max}} = 1.6982$).

**Figure S2: Effect of Flat Damage on Oscillatory Power.**

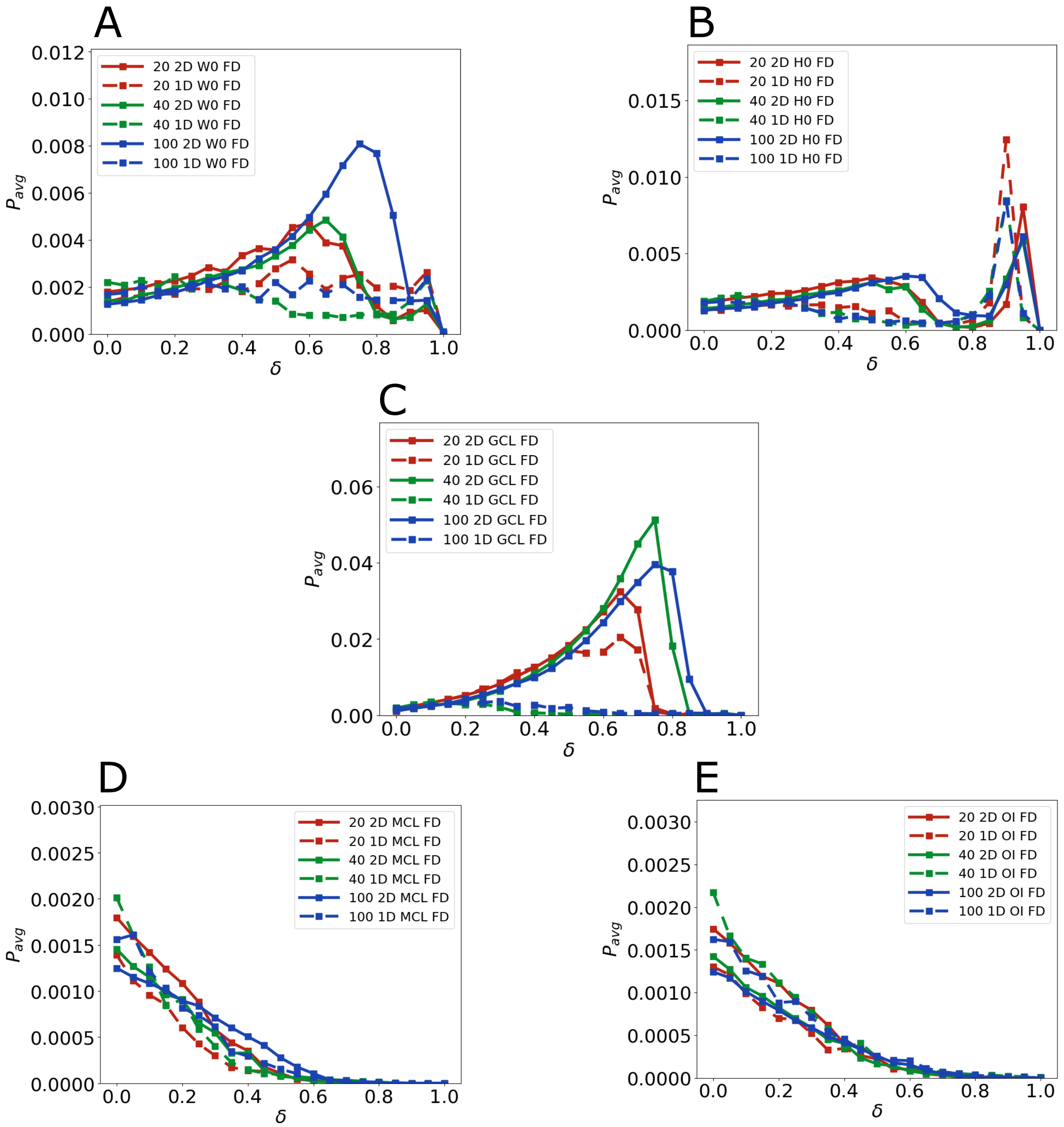


Average ocillatory power ($P_{avg}$) for flat damage to $W_0$ (A), $H_0$ (B), GCL (C), MCL (D), and OI (E) in every network architecture (20 units 1D, 20 units 2D, 40 units 1D, 40 units 2D, 100 units 1D, and 100 units 2D). Solid lines are 2D networks, dashed lines are 1D networks. Note the vertical axes do not have the same scale. The peak in $P_{avg}$ occurred in all 2D networks for flat damage to $W_0$, $H_0$, and GCL.

**Figure S3: Effect of Columnar Damage on Oscillatory Power.**

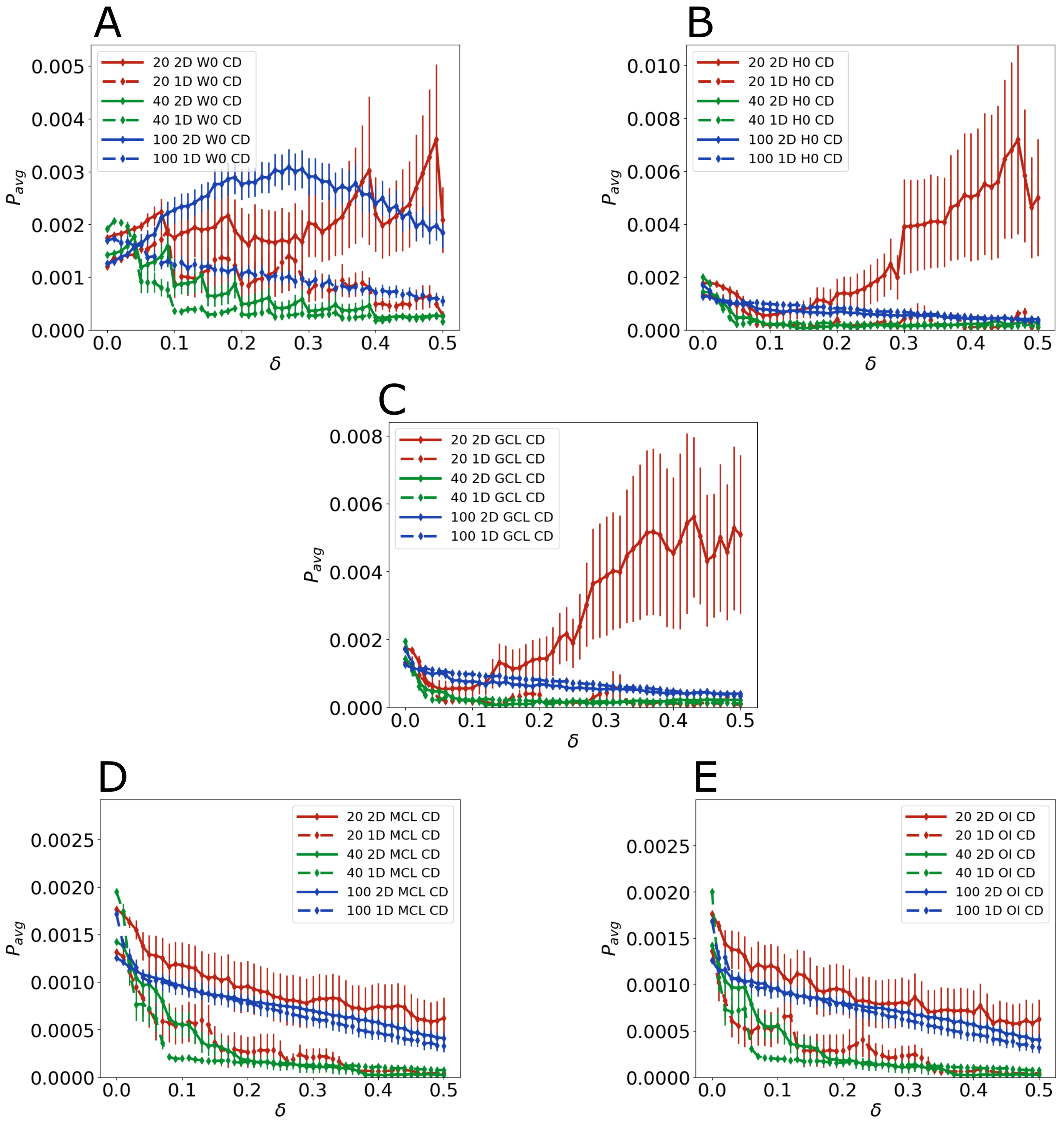


Average oscillatory power ($P_{avg}$) for columnar damage to $W_0$ (A), $H_0$ (B), GCL (C), MCL (D), and OI (E) in every network architecture (20 units 1D, 20 units 2D, 40 units 1D, 40 units 2D, 100 units 1D, and 100 units 2D). Solid lines are 2D networks, dashed lines are 1D networks. Note that the vertical axes do not all have the same scale.

**Figure S4: Effect of Seeded Damage on Oscillatory Power.**

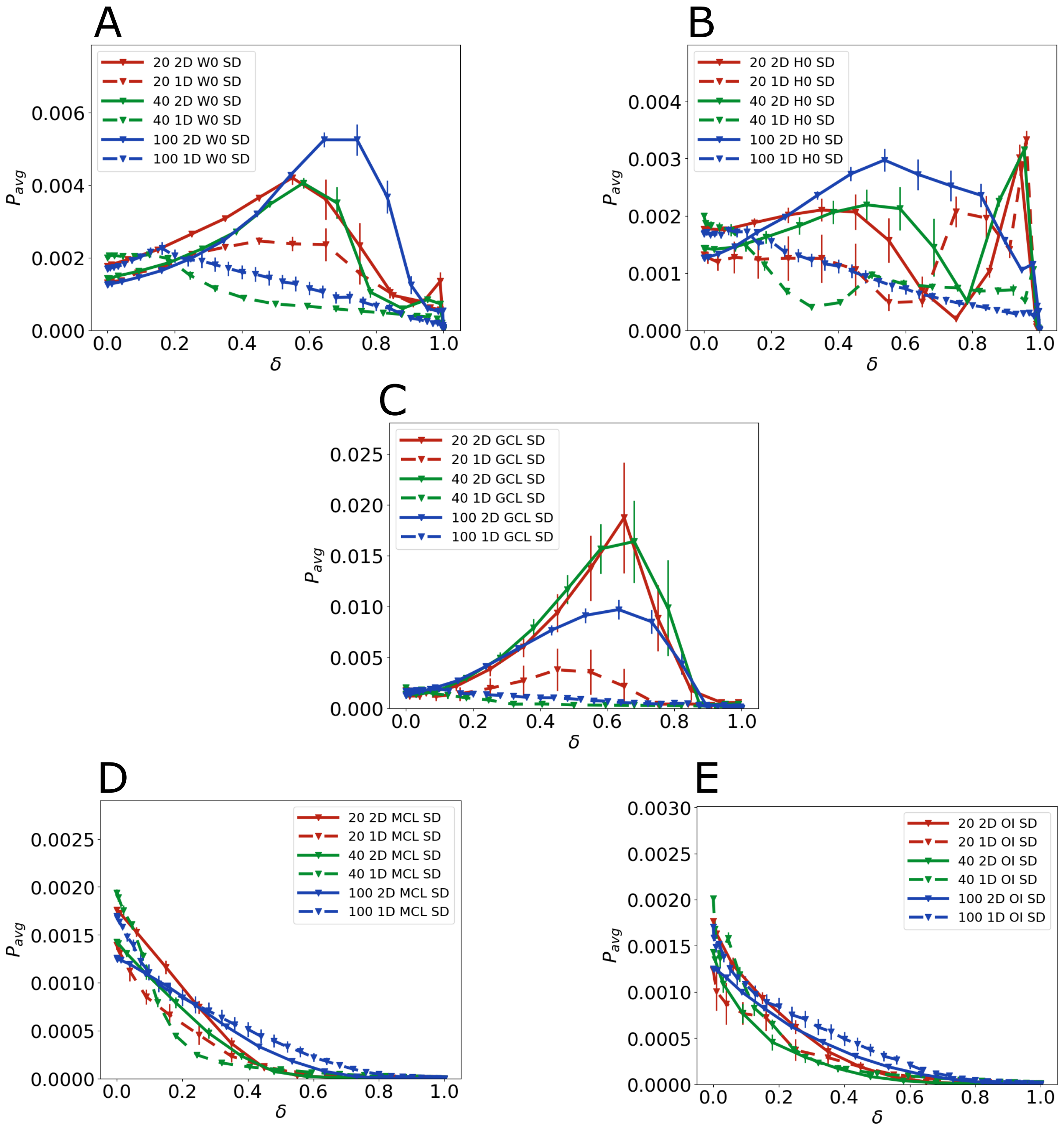


Average oscillatory power ($P_{avg}$) for seeded damage to $W_0$ (A), $H_0$ (B), GCL (C), MCL (D), and OI (E) in every network architecture (20 units 1D, 20 units 2D, 40 units 1D, 40 units 2D, 100 units 1D, and 100 units 2D). Solid lines are 2D networks, dashed lines are 1D networks. Note that not all the vertical axes had the same scale. The peak in $P_{avg}$ occurred in all 2D networks for seeded damage to $W_0$, $H_0$, and GCL.

**Figure S5: Average Oscillatory Power for FD to Both H0 and W0 in the 2D 100 Unit Network**

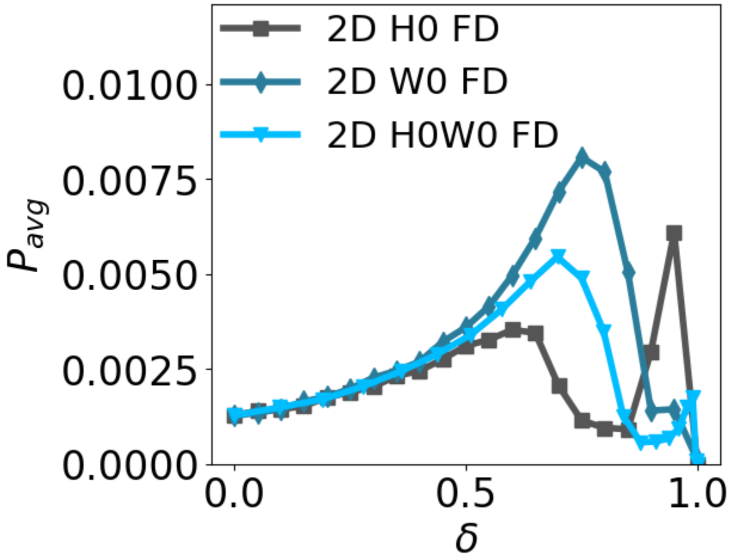


Average oscillatory power for flat damage to both $H_0$ and $W_0$ compared to flat damage to only $H_0$ and flat damage to only $W_0$ in the 2D 100 unit network. Similar to FD to $H_0$ or $W_0$, FD to both resulted in an increase in oscillatory power at intermediate levels of damage. Delivering damage to both seems to result in approximately the mean effect of each individually, in terms of the maximum oscillatory power reached and the location of the maximum.

For this figure, $\delta$ was measured as,

$$\delta = 1 - \frac{\sum_{ij}[H_{\text{Damaged}}W_{\text{Damaged}}]_{ij}}{\sum_{ij}[H_0W_0]_{ij}}.$$

Calculating $\delta$ this way is equivalent to the $\delta$ in the main paper for $W_0$ and $H_0$, as long as the damage delivered is flat damage. Calculating $\delta$ in this manner allows us to compare damage against the same baseline for each target. It is also partly inspired by the linearized analysis summarized in Results, where oscillatory activity was predicted by the matrix,

$$A = H_0 G'_y(\mathbf{y_0}) W_0 G'_x(\mathbf{x_0}),$$

where $G'_y(\mathbf{y_0})$ and $G'_x(\mathbf{x_0})$ are diagonal matrices.

**Figure S6: Evolution of Cell Activity in Seeded Damage to W0 in the 2D 100 Unit Network**

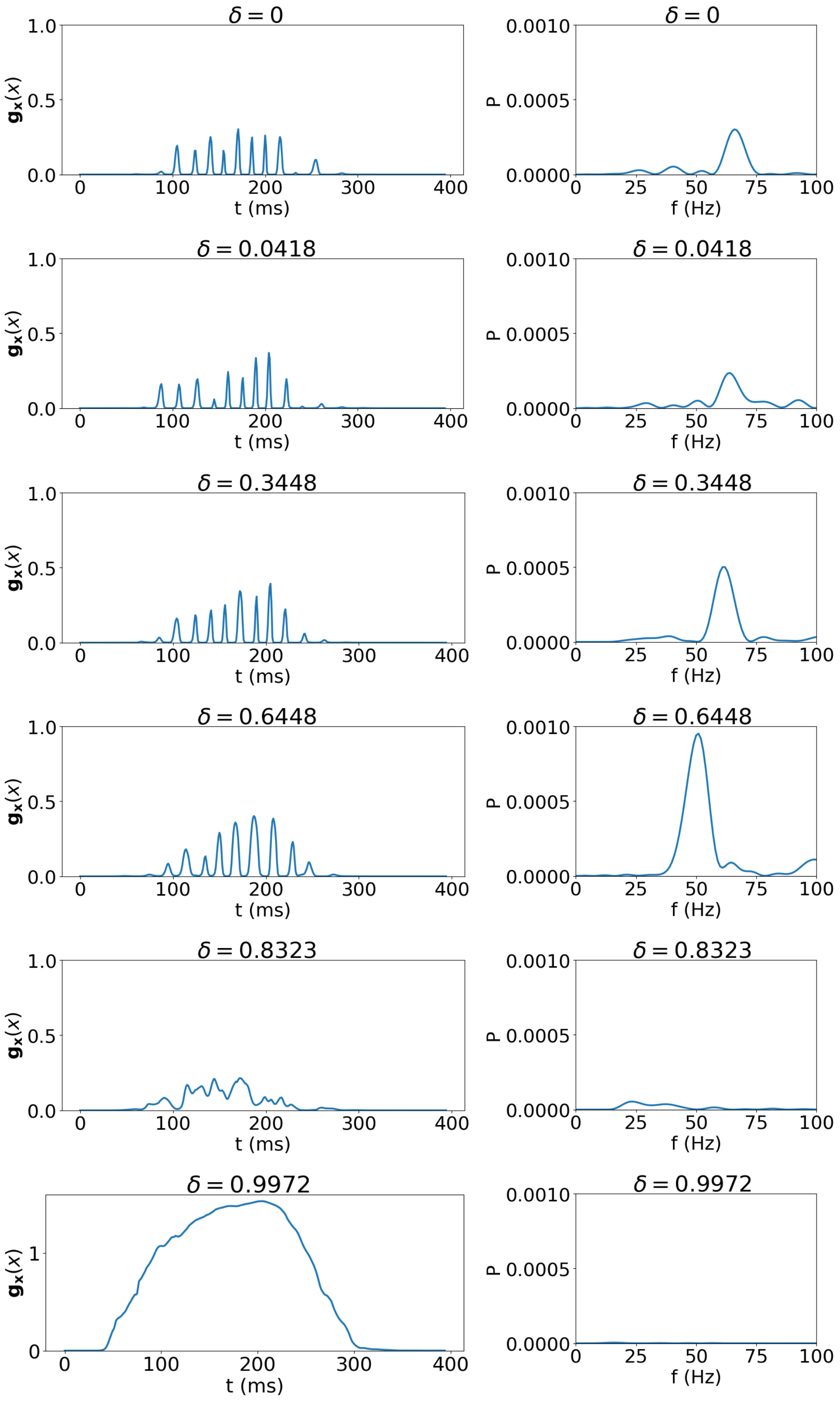


Cell activity ($\mathbf{g_x}$) at various levels of seeded damage to W0 in the 2D 100 unit network, with the associated power spectrum plotted next to each. The inhale peak is at 205 ms. The oscillation amplitude can be seen to grow from $\delta = 0$ to $\delta = 0.8448$, which is reflected clearly in the power spectra. Note that the vertical scale is larger in the last cell activity plot. The sharp fall in activity from about 225 ms to 300 ms can cause an increase in $P_{avg}$, but clearly is not actually an example of gamma band oscillatory behavior and is the reason for the shortened time window used for calculating the power spectra for $P_{avg}$.

As stated in the text, the power spectrum was calculated from the cell activity high-pass filtered above 15 Hz from 125 ms to 250 ms. Some power density was present above 100 Hz, but power density below 100 Hz dominated the contribution to $P_{avg}$.

**Figure S7 Evolution of Cell Activity in Flat Damage to H0 in the 2D 100 Unit Network**

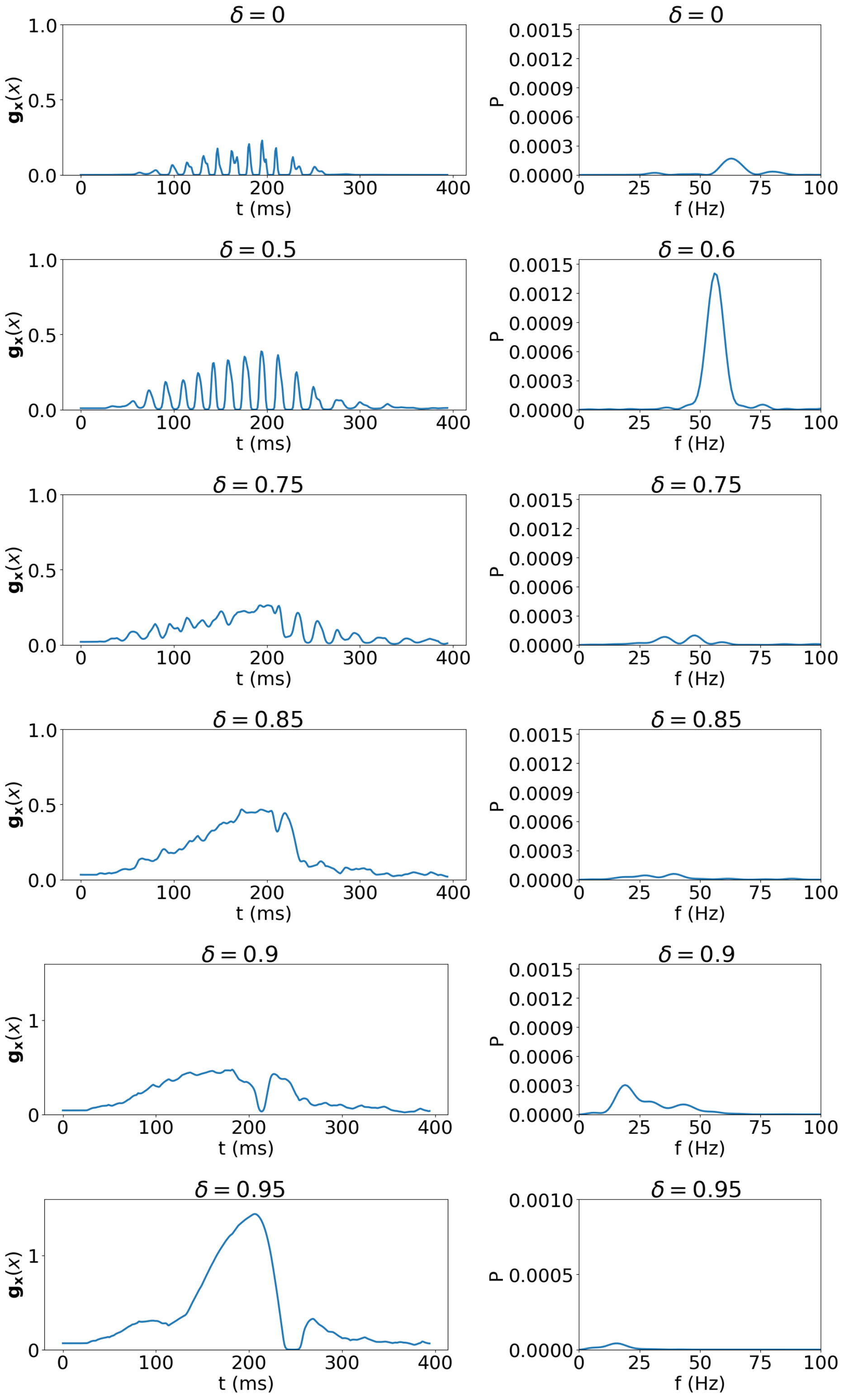


Cell activity ($\mathbf{g_x}$) at various levels of flat damage to $H_0$ in the 1D 100 unit network, with the associated power spectrum plotted next to each. The inhale peak is at 205 ms. The cell activity changes little from $\delta = 0$ to $\delta = 0.5$. Note that the vertical scale is larger in the last two plots of cell activity. The sharp rise then drop seen in $\delta = 0.9$ and $\delta = 0.95$ is the cause of the spike in $P_{avg}$ at the highest levels of damage for flat damage to $H_0$ in the 2D 100 unit network (see Fig. S2B), and is an example of what causes the similar spikes in $P_{avg}$ for flat damage to H0 in all networks (see Fig. S2B and Fig. 4(a) in main text), and for seeded damage to H0 in the 1D 20 unit, 2D 20 unit, and 2D 40 unit networks (see Fig. S4).

As stated in the text, the power spectrum was calculated from the cell activity high-pass filtered above 15 Hz from 125 ms to 250 ms. Some power density was present above 100 Hz, but power density below 100 Hz dominated the contribution to $P_{avg}$.

**Figure S8 Population Activity and Periodograms at Damage Corresponding to Maximum Oscillatory Power for Flat Damage to H0 in 2D 100 Unit Network**

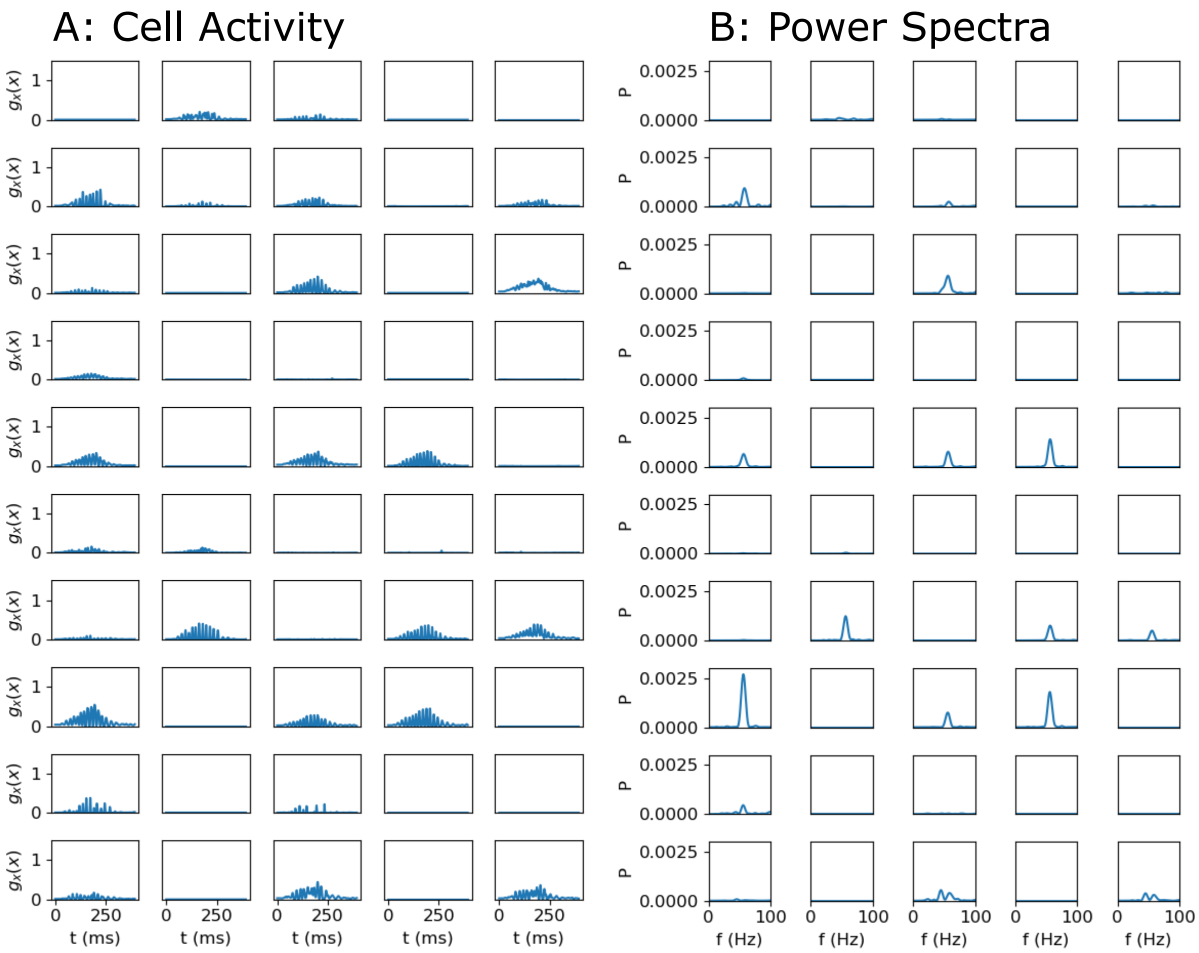


A: Each panel shows the activity of a single MC unit in the 2D 100 unit network at the level of flat damage to $H_0$ corresponding to the greatest amount of oscillatory activity ($\delta = 0.6$). B: Each panel shows the power spectrum corresponding to each MC unit. This demonstrates oscillatory activity giving rise to $P_{\text{avg}}$. Figure S8 demonstrates how spurious $P_{\text{avg}}$ signals can arise at high levels of flat damage to $H_0$.

**Figure S9 Population Activity and Periodograms at Damage Corresponding to Late Peak in Average Power for Flat Damage to H0 in 2D 100 Unit Network**

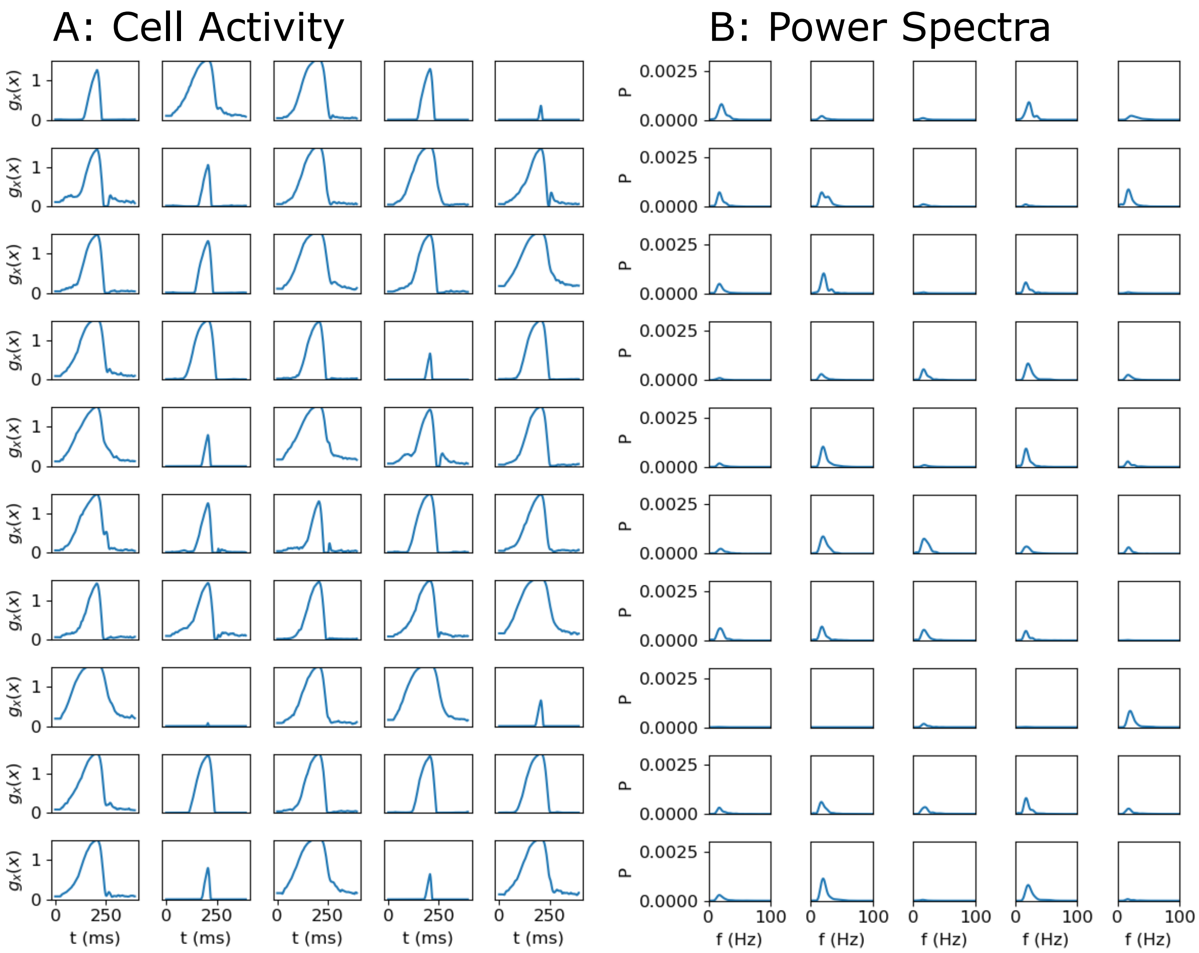


A: Each panel shows the activity of a single MC unit in the 2D 100 unit network at the level of flat damage ($\delta = 0.95$) to $H_0$ corresponding to the sharp peak in $P_{\text{avg}}$ seen in Fig. 4(a) in the main text. B: Each panel shows the power spectrum corresponding to each MC unit. Though most units have a significant signal in their power spectra, no units show oscillations (compare with Figure S8). At high levels of flat damage to H0, the MC unit population reaches higher levels of output state. When odor input decreases, more than half of them drop back down sharply during the time window over which the periodogram is calculated (from 125 to 250 ms). This leads to a large signal in the spectrum and the peak in $P_{\text{avg}}$ that is not actually indicative of oscillatory behavior.
